# SS3D: Sequence similarity in 3D for comparison of protein families

**DOI:** 10.1101/2020.05.27.117127

**Authors:** Igor Lima, Elio A. Cino

## Abstract

Homologous proteins are often compared by pairwise sequence alignment, and structure superposition if the atomic coordinates are available. Unification of sequence and structure data is an important task in structural biology. Here, we present Sequence Similarity 3D (SS3D), a new method for integrating sequence and structure information for comparison of homologous proteins. SS3D quantifies the spatial similarity of residues within a given radius of homologous through-space contacts. The spatial alignments are scored using native BLOSUM and PAM substitution matrices. This work details the SS3D approach and demonstrates its utility through case studies comparing members of several protein families: GPCR, p53, kelch, SUMO, and SARS coronavirus spike protein. We show that SS3D can more clearly highlight biologically important regions of similarity and dissimilarity compared to pairwise sequence alignments or structure superposition alone. SS3D is written in C++, and is available with a manual and tutorial at https://github.com/0x462e41/SS3D/.

## Introduction

Conservation is a term used to describe a protein or nucleic acid sequence that changes slower over time compared to the background mutation rate. For proteins, amino acids that have critical roles in folding, structure, stability, and target recognition tend to be preserved or substituted for ones with similar properties in order to maintain their biological functions (Cooper and Brown, 2008). Conserved regions are often identified through pairwise sequence alignments, structural superposition, and variations thereof (Doğan and Karaçalı, 2013). Sequence alignments involve the linear comparison of two or more amino acid sequences, and are typically created using computer algorithms that attempt to maximize a scoring function based on a substitution matrix in which aligned identical positions are scored most favorably, followed by alignment of amino acids with similar biochemical properties. The alignment of dissimilar amino acids, or introduction of gaps negatively impacts the alignment score. In cases where evolutionary relatedness is difficult to ascertain from a sequence alignment, structural superposition can be useful. The degree of structural similarity between two proteins is commonly quantified using the root mean squared deviation (RMSD) of the atomic coordinates, where lower values indicate a higher degree of 3D similarity.

Sequence alignments and structural superposition have their own benefits and downsides. While a sequence alignment allows for quantification of conservation at every position, in the absence of accompanying structures, there is no knowledge of the 3D organization of the linear sequence. As a result, conserved regions, such as active sites and interaction surfaces, which are commonly formed by the association of residues distant in primary sequence, can be difficult to identify, characterize, and compare. On the other hand, superpositions of protein structure allow for visual comparison of such regions and quantification of geometrical similarity, often using global or local RMSD (Maiti et al., 2004). The approach also has limitations. For instance, functionally different proteins can share similar folds, leading to a close superposition, which may lead one to incorrect assumptions regarding function, substrate specificity, or other properties.

To overcome some of the limitations intrinsic to pairwise sequence alignments and structural superposition, we have developed a unique method, dubbed SS3D, for comparing homologous proteins using both sequence and structure information. In the following sections, we describe the SS3D method, and use several case studies to demonstrate its ability to better distinguish biologically important regions of similarity and dissimilarity compared to pairwise sequence alignment or structure superposition alone.

## Methods

### Requirements and program execution

To compare two homologous proteins using SS3D, all that is required is a structure file in pdb format of each molecule. The atomic coordinates may be obtained from the protein data bank, model databases, or modeling approach of choice. It is not necessary to superimpose the structures, as SS3D analysis utilizes residue numbers, which must be the same for homologous positions in the two proteins. A simple tutorial for accomplishing this using Chimera (Pettersen et al., 2004) is included with the tool. First, the software builds a Cα-Cα contact map of each protein. By default, a 10 Å Cα-Cα cutoff distance is used, but the parameter is adjustable by the user. For each homologous Cα-Cα contact between the two proteins, a list of residues that lie within a user specified 3D search radius of either Cα atom involved in the contact (spatial neighbor list) is obtained separately for each protein (Fig. 1a,b). The spatial alignment is then scored using the standard BLOSUM62 or PAM250 substitution matrices (Fig. 1b); smaller values indicate a lower similarity, and larger values correspond to a higher similarity around the contact. The program contains additional user adjustable options for refining residue selection, output of the individual alignments, and normalization of the scores on a 0-1 scale. All examples shown here were executed with default code parameters.

**Fig. 1.**
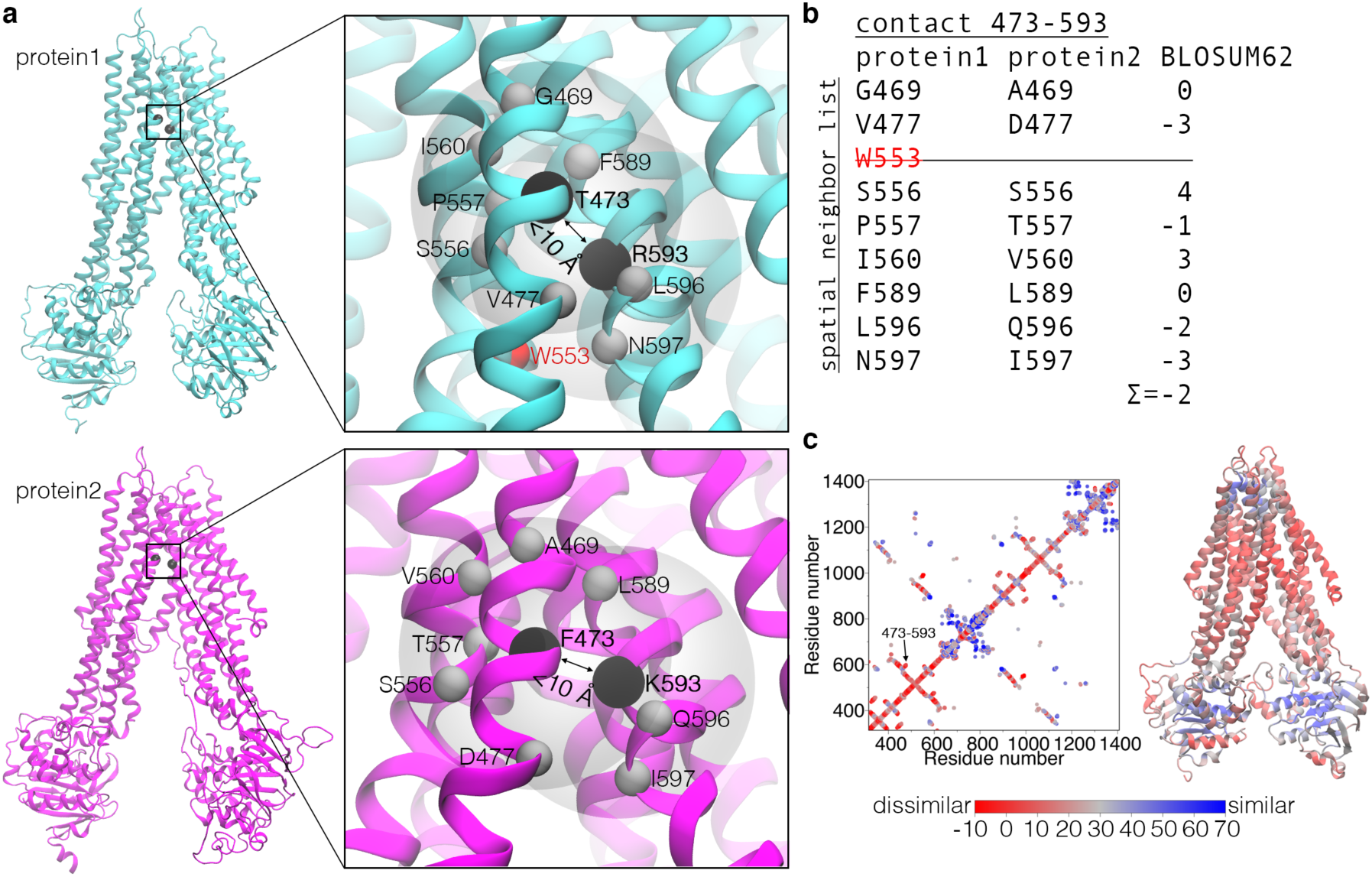
Depiction of the SS3D method. a) Example of a homologous Cα-Cα contact (black spheres) in two proteins of the same family, and Cα atoms within the user-specified 3D search radius of either Cα atom involved in the contact. b) Spatial alignment and scoring of homologous positions in the vicinity of the contact. Note that the three residues directly preceding and succeeding the ones involved in the contact are not included in the analysis, despite falling within the search radius. Exclusion of neighboring positions is user adjustable, and can be useful to decrease the contribution of sequential residues. Also note, W553 of protein1 is not included in the scoring because the homologous position in protein2 falls slightly outside of the search radius. c) Matrix and B-factor representations of the SS3D result.

### Data representation

The primary SS3D results are output in two convenient formats. The standard output is a text file containing the residue numbers of the Cα atoms forming a contact, followed by their spatial similarity score, one per line (e.x. 473 593 -2) (Fig. 1b). This output can be visualized using a contact matrix colored by score (Fig. 1c). SS3D can also map spatial similarity scores to the B-factor column of the input pdb files, which can be colored by value in protein structure visualization software (Fig. 1c). In this work, we used a red-gray-blue spectrum to represent low, medium, and high spatial similarity scores, respectively. This representation is effective for illustrating the correlation between similarity and protein structure-activity relationships, as demonstrated herein.

## Results and discussion

To demonstrate SS3D, members of five different protein families were assessed: GPCR, p53, kelch, SUMO, and SARS coronavirus spike protein. For each case, two family members were compared by pairwise sequence alignment scored with BLOSUM62 (linear similarity), structure superposition (RMSD), and spatial similarity scored with BLOSUM62 (SS3D). In doing so, we demonstrate that SS3D can more clearly highlight physiologically important regions of similarity and dissimilarity than either method alone. We expect that SS3D will be useful in numerous applications, including, but not limited to, improved detection of conserved and nonconserved regions, delineation of evolutionary pathways, assessment of interaction specificity, and protein design. The following cases illustrate SS3D utility for these purposes.

### Case 1: GPCR

The GPCR superfamily is the largest class of mammalian receptors (Baltoumas et al., 2013). GPCR family members share the greatest homology in the conserved transmembrane (TM) segments, and highest diversity around the extracellular amino terminus (Kobilka, 2007). There are over 800 GPCRs in humans that mediate a diverse array of physiological processes such as taste, vision, and smell by responding to extracellular stimuli, including peptides, ions, small molecules, and photons. Figure 2a-c shows the pairwise, structural, and SS3D comparisons of two class A GPCRs, β_2_ adrenergic receptor (β_2_AR), and type 1 cannabinoid receptor (CB1). Among the comparisons, SS3D reveals the different target specificities of the two receptors most apparently (Fig. 2c). The primary endogenous agonists of β_2_AR are epinephrine and norepinephrine (Reiner et al., 2010), while that of CB1 is anandamide (Fig.2d) (Felder and Glass, 1998). β_2_AR and CB1 agonists bind to a pocket formed by TM segments in the extracellular leaflet, and the residues that make physical contacts with these ligands (Chan et al., 2016; Shao et al., 2016) present below average SS3D scores (Fig. 2c,e-f). In contrast, the TM segments in the intracellular leaflet have higher SS3D values, specifically around the (D/E)RY and NPXXY segments (Fig. 2c), which have critical roles in global movements of the helices required for G protein binding and activation, which is common among class A GPCRs (Katritch et al., 2013).

**Fig. 2.**
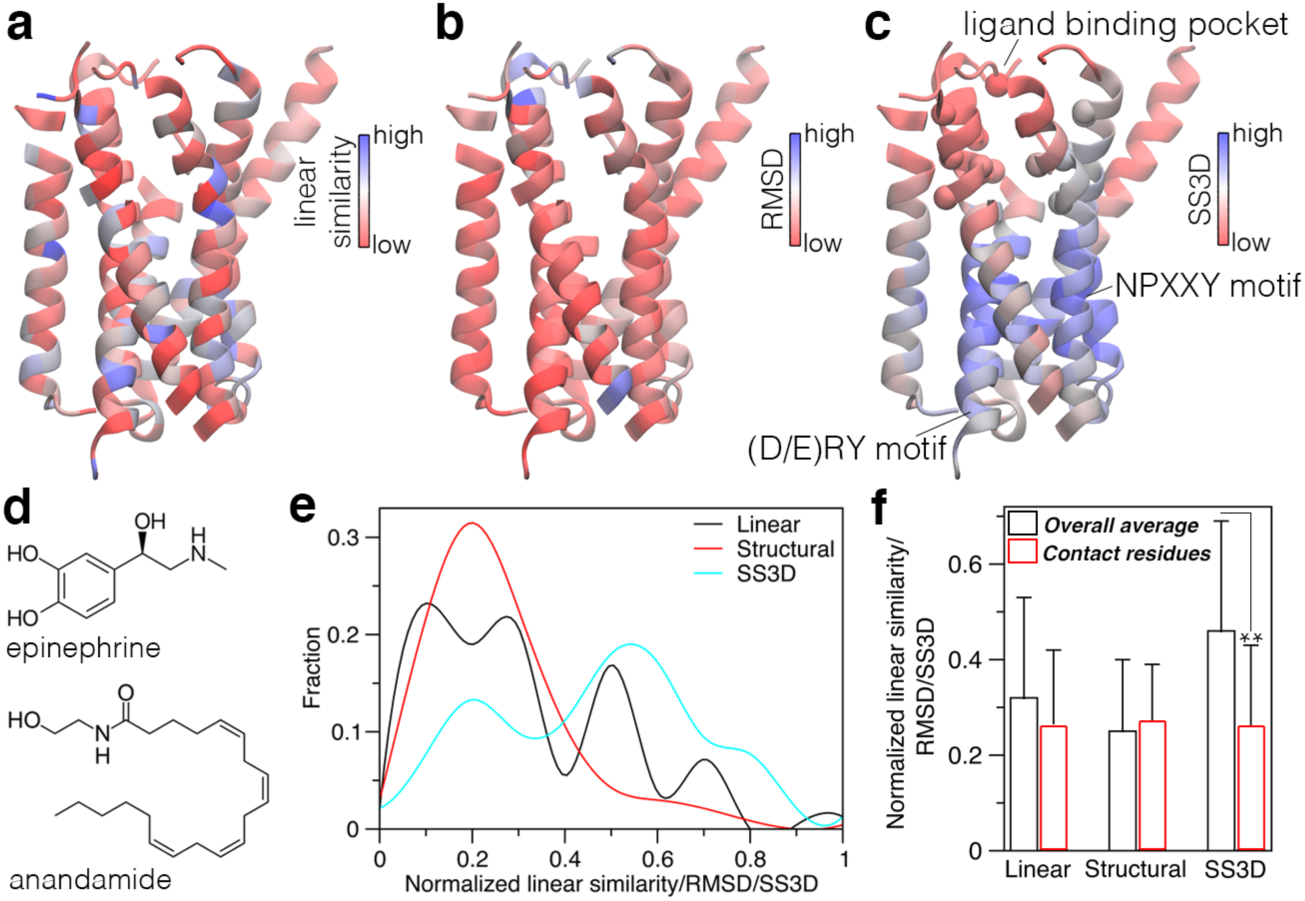
Comparison of β_2_AR and CB1. Pairwise sequence comparison of β_2_AR and CB1 mapped onto the β_2_AR structure, 2rh1 (Cherezov et al., 2007). b) Cα RMSD values between β_2_AR and CB1 mapped on the β_2_AR structure. c) SS3D comparison of β_2_AR, and CB1, 5u09 (Shao et al., 2016) showing nonidentical residues that make physical contacts with β_2_AR or CB1 ligands as spheres (M36, V39, V87, A92, W109, T110, D113, V114, F193, S203, S204, F289, F290, V292, N293, Y308, I309, N312; β_2_AR residue type and numbering used). d) Chemical structures of the main endogenous ligands of β_2_AR and CB1. e) Distribution of the normalized pairwise, RMSD, and SS3D values. f) Comparison of the overall averages, and averages of residues that make physical contacts with β_2_AR ligands (** t-test *p*<0.005).

### Case 2: p53

The p53 family of tumor suppressors contains three members, p53, p63, and p73, which have both similar and distinct functions. The DNA binding domains (DBDs) of these proteins are the most conserved regions, sharing ∼60% sequence similarity and analogous structures (Costa et al., 2016). Physical contact with DNA occurs at the loops protruding on the top portion of the β-sandwich fold (Fig. 3). p53 family members bind to similar response elements, and thus have overlapping transcriptional profiles (Ciribilli et al., 2013). The high conservation of the DNA binding loops is considerably less evident in the pairwise alignment and structure superposition (Fig. 3a,b) compared to SS3D (Fig. 3c). Specifically, DNA contact residues N239, Q136, R273, and R248W (Freed-Pastor and Prives, 2012) show among the highest conservation levels of the entire DBD in the SS3D comparison with an average similarity score of 0.91 (Fig. 3c). The same residues have an average score of 0.68 in the pairwise alignment (Fig. 3a).

**Fig. 3.**
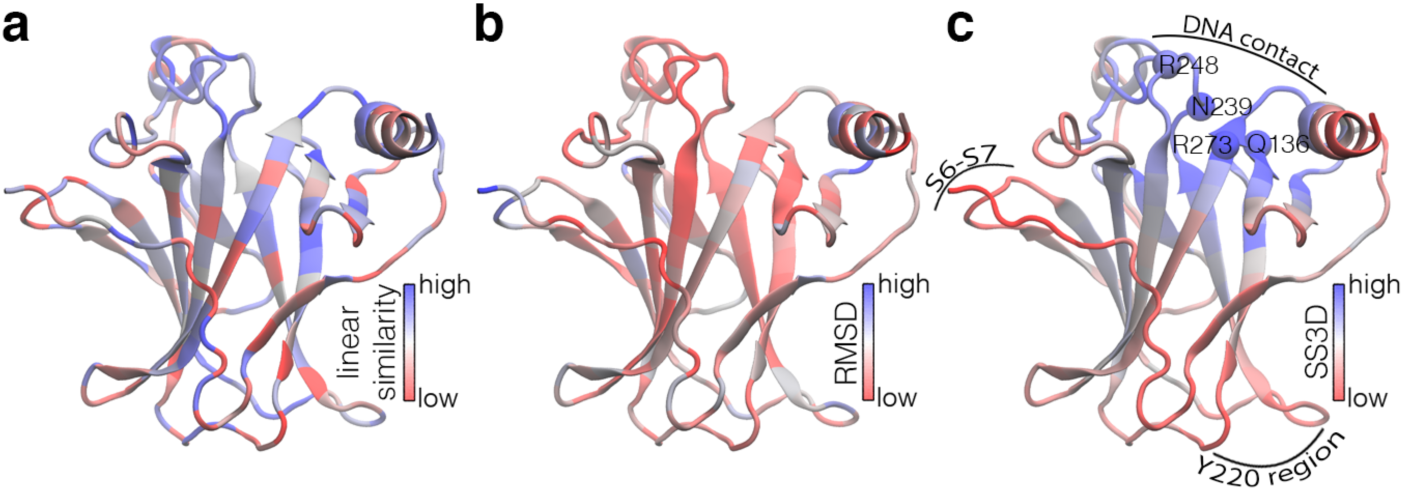
p53 and p63 DBD comparison. a) Pairwise comparison of p53 and p63 DBD sequences mapped to the p63 DBD structure, 2rmn (Enthart et al., 2016). b) Cα RMSD values between p53 and p63 DBDs mapped onto the p63 DBD structure. c) SS3D comparison of p53, 2xwr (Natan et al., 2011), and p63 DBDs.

In addition to accentuating regions of high similarity, SS3D also conveniently illustrates differences. Despite their high sequence and structure similarity, p53 family members vary considerably in stability, with p53 being the least stable and most aggregation prone family member (Cino et al., 2016). p63 is the family member with highest stability and lowest tendency to aggregate. The S6-S7 and Y220 regions (Fig. 3c) exhibit altered conformational sampling in p53 and structurally compromised mutants, leading to formation of surface crevices that destabilize the p53 DBD, promoting its unfolding and aggregation (Pradhan et al., 2019; Xu et al., 2012). Both pockets are being explored as targets for p53 stabilizing molecules (Li et al., 2019). The SS3D comparison of p53 and p63 DBDs shows low similarity in these areas (Fig 3c). Therefore, the SS3D analysis could potentially guide the design of higher stability variants, which is another proposed strategy for restoring p53 function (Lima et al., 2020).

### Case 3: kelch domain

The kelch repeat motif is a 44-56 amino acid sequence that forms a four-stranded antiparallel beta sheet (Adams et al., 2000). The repeats, or blades, can associate, typically in clusters of 4-7, to form β-propellers. Kelch repeat proteins are involved in diverse cellular functions, and interact with binding partners through residues surrounding a central cavity (Fig. 4). The kelch domain of KLHL2 selectively recognizes ‘EXDQ’ motifs (Gouw et al., 2018), while that of KLHL19 binds to ‘EXGE’ containing motifs (Karttunen et al., 2018). Both have high selectivity for their particular consensus motifs, and are unable to bind to those of one another interchangeably (Schumacher et al., 2014). The SS3D comparison of KLHL2 and KLHL19 kelch domains (Fig. 4c) shows a discernable pattern, where blades 1 and 2 have lower similarity scores relative to blades 3-6, which is not evident by pairwise alignment or RMSD (Fig. 4d,e). Intriguingly, crucial residues for KLHL19 interaction with its major partner, nrf2, are located within blades 1 and 2 (Karttunen et al., 2018): Y334 (blade 1), R380 (blade 2), and N382 (blade 2). Because KLHL2 and KLHL19 share a common ancestor, it seems plausible that individual blades evolved at different rates, and greater divergence of a subset of the repeats (blades 1 and 2) was sufficient to achieve novel interaction specificity.

**Fig. 4.**
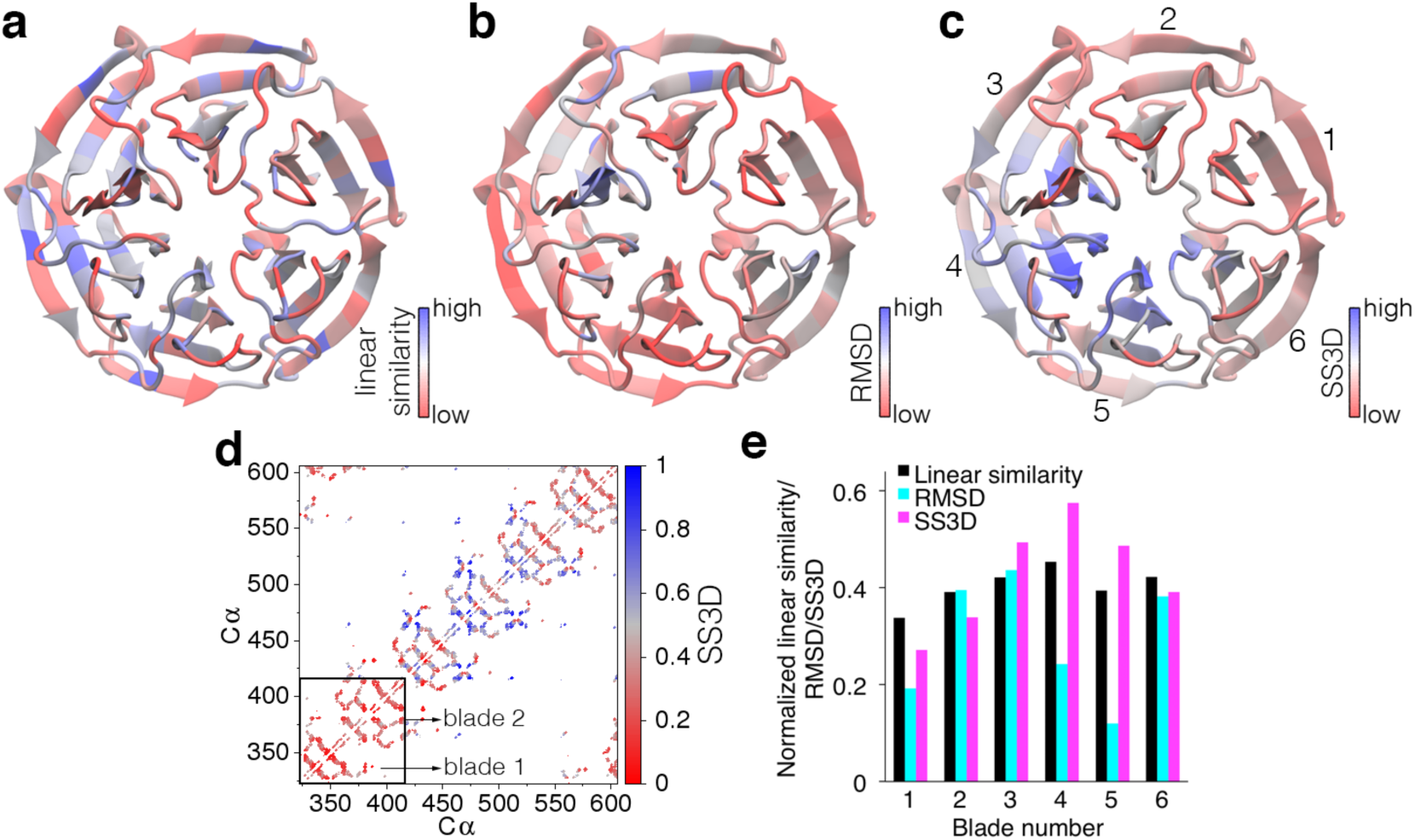
Comparison of KLHL2 and KLHL19 kelch domains. a) Pairwise comparison of the KLHL2 and KLHL19 kelch domain sequences, mapped onto the KLHL19 structure, 1u6d (Li et al., 2004). b) Cα RMSD values between KLHL2 and KLHL19 kelch domains mapped onto the KLHL19 structure. c) SS3D comparison of KLHL2, 2xn4 (Canning et al., 2013), and KLHL19, 1u6d (Li et al., 2004). d) SS3D matrix representation. e) Linear, RMSD, and SS3D comparisons of the individual blades.

### Case 4: SUMO

Ubiquitin and ubiquitin-like proteins originate from a common β-grasp fold ancestor, and comprise a diverse family with numerous cellular functions (Zuin et al., 2014). Differently from ubiquitin, which targets proteins for degradation, small ubiquitin-like modifiers (SUMOs) are key regulators of transcription, apoptosis, cell cycle, and other fundamental processes (Müller et al., 2001). SUMOs can bind both covalently and noncovalently to target proteins. The former is catalyzed by SUMO-specific proteases (SENPs), while the latter is mediated by SUMO interacting motifs (SIMs) (Hickey et al., 2012). SENPs and SIMs have been shown to have SUMO isoform preference, with SUMO1 residues Q53, R54, A72, H75, and G81, being important determinants of interaction specificity (Chupreta et al., 2005; Ronau et al., 2016). Compared to the pairwise alignment and structure superposition of SUMO1 and 2 (Fig. 5a,b), SS3D better highlights the differences around these residues, which present low spatial similarity scores (Fig. 5c). The comparison also illustrates the power of SS3D in identifying conserved regions. Although the ubiquitin-like protein family has become highly diversified, the ancestral β-grasp scaffold has been preserved. SS3D precisely shows high conservation around the protein core, and even reveals differential conservation of surface- and core-oriented residues (Fig. 5c).

**Fig. 5.**
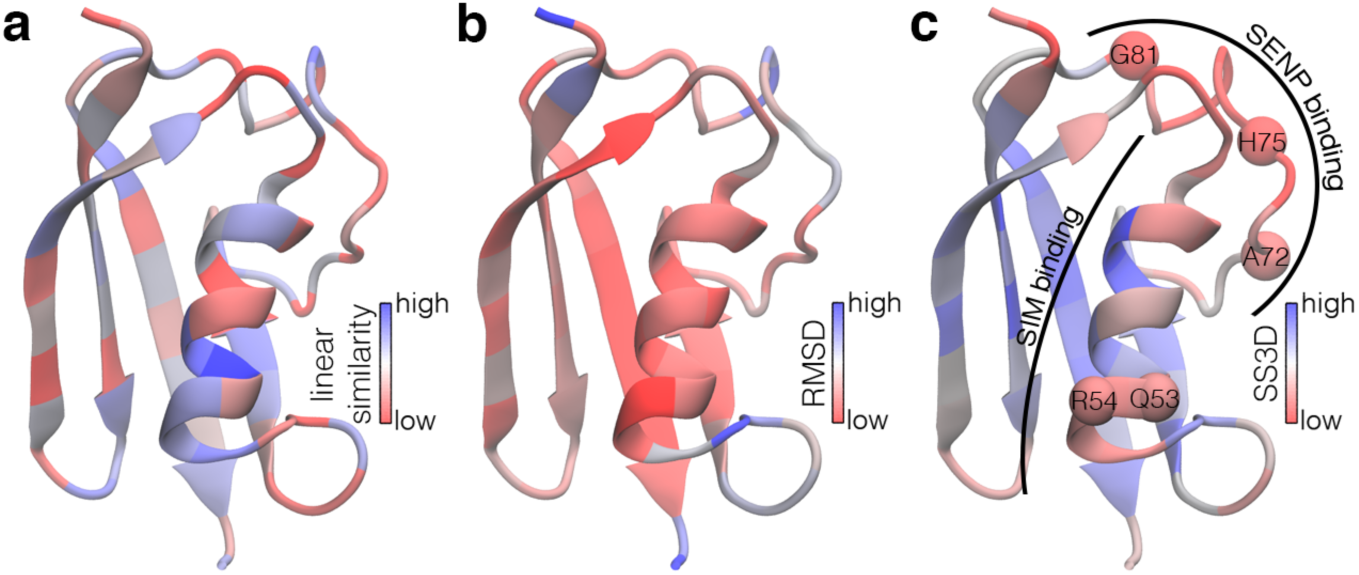
Comparison SUMO1 and SUMO2. a) Pairwise comparison of SUMO1 and 2 sequences mapped onto the SUMO1 structure, 1a5r (Bayer et al., 1998). b) Cα RMSD values between SUMO1 and 2 mapped onto the SUMO1 structure. c) SS3D comparison of SUMO1 and 2, 2n1w, with important residues for SENP and SIM interaction specificity drawn as spheres.

### Case 5: SARS coronavirus spike protein

The spike protein is a large glycosylated homotrimer that garnishes the viral envelope and interacts with host receptors permitting virus fusion with cell membranes (Belouzard et al., 2012). Spike protein neutralizing antibodies have been detected in coronavirus patients, implicating it as a prime antigenic component and target for vaccine development (Kitabatake et al., 2007). Although the SARS-CoV-1 and SARS-CoV-2 spike proteins share 76% sequence identity (Jaimes et al., 2020), several antibodies against the former spike protein do not cross-neutralize that of the latter, indicating that they possess unique antigenic characteristics (Yi et al., 2020). Assessment of the differences between key features in spike protein variants is expected to be useful for development of vaccines and virus entry inhibitors (Robson, 2020). The N-terminal and receptor binding domains (NBD and RBD) are potential targets for neutralizing antibodies (Jiang et al., 2020), while the S1/S2 cleavage site facilitates virus entry and has been determined essential for SARS-CoV-2 infection of humans (Hoffmann et al., 2020). Interestingly, these regions are more prominently seen by SS3D comparison than pairwise or structural similarity alone (Fig. 6a-d). Within the RBD, the ACE2 receptor binding motif (RBM) is a crucial target for neutralizing antibodies (Freund et al., 2015). SS3D reveals low similarity in the central portion of the RBM (Fig. 6e), which could explain poor cross-neutralizing activity of most SARS-CoV-1 antibodies against SARS-CoV-2 (Lan et al., 2020). Neither the pairwise similarity nor RMSD comparisons are able to discriminate RBM uniqueness (Fig. 6f).

**Fig. 6.**
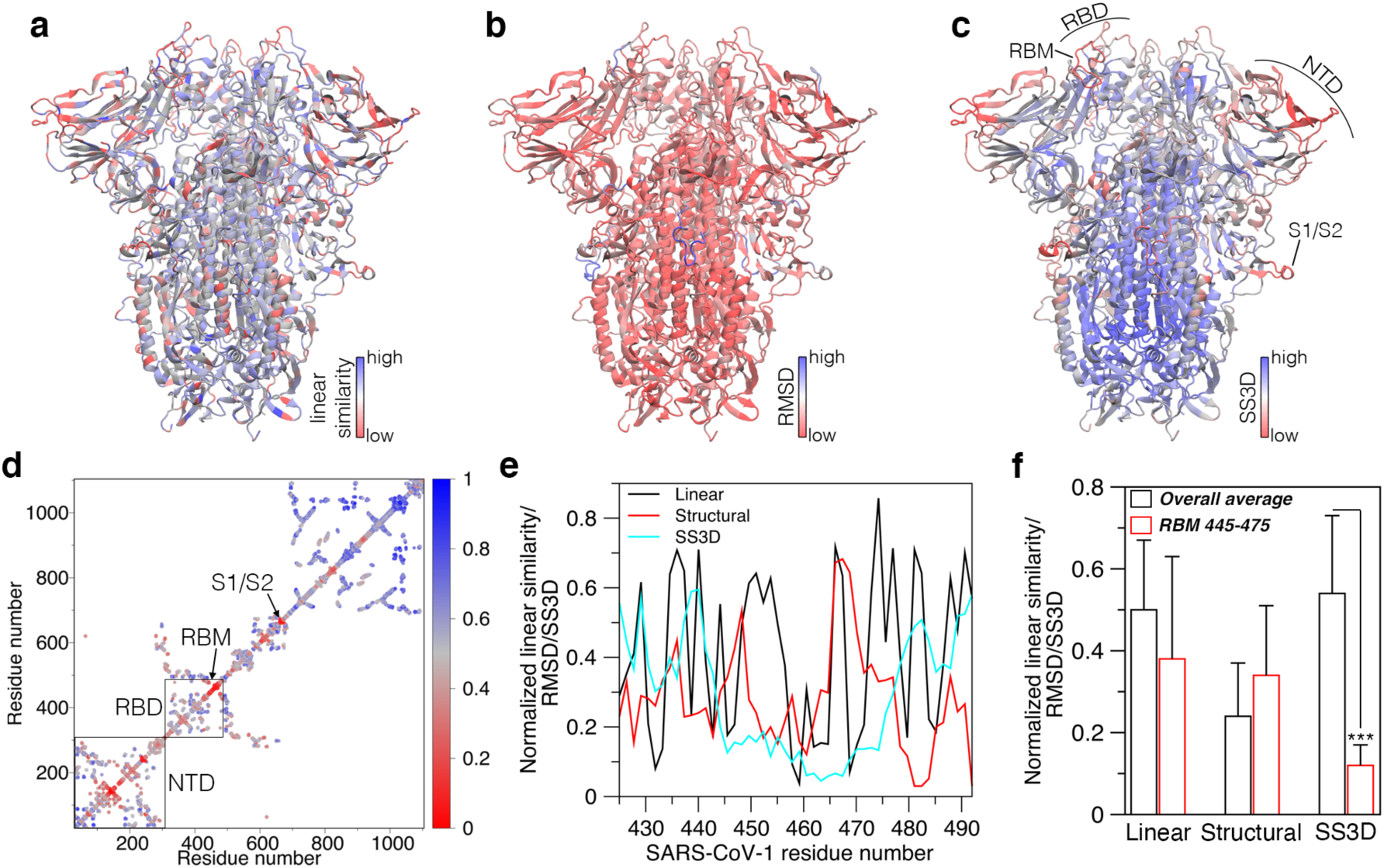
Comparison of SARS-CoV-1 and SARS-CoV-2 spike proteins. a) Pairwise comparison mapped onto the SARS-CoV-1 prefusion spike protein structure, 5×58 (Yuan et al., 2017). b) Cα RMSD values between SARS-CoV-1 and SARS-CoV-2 spike protein, 6vxx (Walls et al., 2020) mapped onto the SARS-CoV-1 structure. c) SS3D comparison of the two spike proteins. d) SS3D matrix representation of the monomeric unit. e) Normalized pairwise, RMSD, and SS3D values of the RBM. f) Comparison of the overall averages, and averages of core RBM residues 445-475 (SARS-CoV-1 numbering; *** t-test *p*<0.001).

## Conclusions

The present work introduces a new method for comparing homologous proteins by integrating sequence and structure information. By employing a standard scoring metric, SS3D provides a quantitative measure of spatial similarity between proteins. Case studies of different protein families illustrate that SS3D more effectively shows biologically important regions of similarity and dissimilarity compared to pairwise sequence alignments or structure superposition alone. We anticipate that the tool will be useful for numerous bioinformatics purposes including, and beyond those illustrated here.

## Notes

### Competing Interest Statement

The authors have declared no competing interest.

